# Antifungal susceptibility testing of *Aspergillus niger* on silicon microwells by intensity-based reflectometric interference spectroscopy

**DOI:** 10.1101/804385

**Authors:** Christopher Heuer, Heidi Leonard, Nadav Nitzan, Ariella Lavy-Alperovitch, Naama Massad-Ivanir, Ester Segal

## Abstract

The increasing number of invasive fungal infections among immunocompromised patients and the emergence of antifungal resistant pathogens has resulted in the need for rapid and reliable antifungal susceptibility testing (AFST). Accelerating antifungal susceptibility testing allows for advanced treatment decisions and the reduction in future instances of antifungal resistance. In this work, we demonstrate the application of a silicon phase grating as sensor for the detection of growth of *Aspergillus niger* (*A. niger*) by intensity-based reflectometric interference spectroscopy and its use as an antifungal susceptibility test. The silicon gratings provide a solid-liquid interface to capture micron-sized *Aspergillus* conidia within microwell arrays. Fungal growth is optically tracked and detected by the reduction in the intensity of reflected light from the silicon grating. The growth of *A. niger* in the presence of various concentrations of the antifungal agents voriconazole and amphotericin B is investigated by intensity-based reflectometric interference spectroscopy and used for the determination of the minimal inhibitory concentrations (MIC), which are compared to standard broth microdilution testing. This assay allows for expedited detection of fungal growth and provides a label-free alternative to standard antifungal susceptibility testing methods, such as broth microdilution and agar diffusion methods.

## Introduction

Pathogenic fungi are a rising cause of disease and an emerging threat among immunocompromised individuals.^1^ In particular, *Candida albicans* and pathogenic *Aspergilli*, such as *A. fumigatus*, account for the majority of invasive fungal infections,^2,3^ resulting in an estimated 1.4 million deaths by fungal infections worldwide every year.^4^ Because of acquired antimicrobial resistance, specifically antifungal resistance in yeast and filamentous fungi, species identification alone is not sufficient enough to target efficient therapy for fungal infections, emphasizing the need for a reproducible and rapid antifungal susceptibility testing (AFST).^5,6^ Thus, improving and accelerating AFST will abate the emergence of antifungal resistance and reduce the number of deaths associated with ineffective prescription of an antifungal therapy.^1,3,7^

Classically employed methods for AFST include broth microdilution (BMD) testing as the suggested reference method standardized by the European Committee on Antimicrobial Susceptibility Testing (EUCAST) and the Clinical and Laboratory Standard Institute, as well as agar diffusion methods, such as the commercially available Etest (bioMérieux SA). However, these methods are time-consuming (MIC determination after 24 - 72 h) and not intended for routine use.^8^ Automated and commercially available BMD tests such as the Vitek2 (bioMériuex SA) and the Sensititre YeastOne (Thermo Fisher) are easy to use, but cost-intensive and limited to antifungal susceptibility testing of yeast.^9^

New approaches for molecular identification of fungal pathogens and rapid AFST include nucleic acid based diagnostics and mass spectrometry.^8,10,11^ In particular, matrix-assisted laser desorption-ionization time of flight mass spectrometry (MALDI-TOF MS) has been used for species discrimination and the identification of resistance by analyzing protein fingerprints of fungal isolates.^12–14^ Rapid AFST by MALDI-TOF MS is achieved within 3 h by investigating proteomic changes due to exposure of fungal cells to antifungal agents.^14,15^ Nonetheless, the disadvantages of mass spectrometry for AFST include high acquisition costs, the need for optimized preparation protocols and further validation, as well as the ability to detect resistance only in selected cases.^6,12^

Genotypic AFST consisting of nucleic acid-based strategies to identify resistance-conferring mutations^8^ can be expensive^12^ and limited due to the lack of identifiable resistance markers for some antifungal agents and multiple mutations and mechanisms that can confer resistance.^11,16,17^ Furthermore, genomic AFST does not allow for phenotype-based determination of minimal inhibitory concentrations (MIC) and is not yet standardized for practical use in most clinical laboratories,^11^ suggesting that the development of rapid, sensitive and affordable phenotypic methodologies for AFST is crucial. Therefore, in recent years, research has focused on the development of novel methods for phenotypic antifungal susceptibility testing such as flow cytometry^18,19^, colorimetric redox indicators^20–22^, isothermal microcalorimetry^23,24^, and porous aluminum oxide assays^5^. Furthermore, microfluidic systems for single cell susceptibility testing and diagnostic applications have been developed recently. ^25–27^ Many of these new techniques rely on costly equipment, have a limited microorganism spectrum or do not yet effectively accelerate the determination of MIC values.

The aim of this work is to develop an easy-to-perform, affordable platform that allows the growth of *Aspergillus* species to be tracked in real time, enabling rapid AFST. This platform is based on the optical sensing of reflected light from a micropatterned diffraction grating.^28^ Previously, phase-shift reflectometric interference spectroscopic measurements (PRISM) was demonstrated to monitor antibiotic susceptibility and the behavior of bacteria within microstructured arrays.^29–33^ Reflectance spectra of the arrays were collected over time in order to infer values of *2nL*, in which *n* represents refractive index of the medium within the arrays and *L* represents the height of the microstructures. However, due to the different behavior and morphology of filamentous fungi compared to bacteria, herein we apply intensity-based PRISM, referred to as iPRISM, as a principle for the detection of microorganisms and as a tool for label-free, phenotypic antifungal susceptibility testing using fungal species *A. niger* as a model microorganism.

## Results and Discussion

### Principals of the iPRISM Assay for AFST

iPRISM for antifungal susceptibility testing relies on the capture of fungal conidia within silicon microwells and the subsequent monitoring of fungal growth in real time by intensity-based reflectometric interference spectroscopy measurements, as depicted in Figure 1. Assays are performed in a series of temperature-controlled microfluidic channels each containing a syringe injection port and an outlet to waste. Photonic chips consisting of Si diffraction gratings, specifically periodic microwell arrays with a width of ~3 µm and a depth of ~4 µm, are individually fixed in the center of the flow channels and illuminated by a tungsten-halogen white light source (LS-1, Ocean Optics) positioned normal to the Si chip (Fig. 1A). The resulting reflectance spectrum of the zero-order diffraction exhibits interference fringes (Fig. 1B), as the incident light is partially reflected by the top and the bottom of the microwells.^29,34^ Applying frequency analysis results in a single peak where the peak position corresponds to the *2nL* and the peak amplitude or intensity (*I*) corresponds to the intensity of the reflected light (Fig. 1D-i and 1D-ii). Whereas in previous works^28–33^ the values of *2nL* were monitored over time, in this assay, the intensity of the reflected light is tracked, as *A. niger* tends to grow on top of the silicon microwells resulting in a decrease in peak intensity. The percent change in peak intensity of the fast Fourier spectrum, *ΔI*, of the reflected light over time is calculated as:

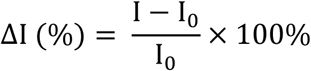

in which *I* is the intensity value at a given time and *I*_*0*_ is the first intensity value after the introduction of the conidia suspension during growth monitoring experiments or when antifungal was first introduced into the system during AFST experiments (deemed time 0). iPRISM for AFST of *A. niger* is performed in two steps: Initially, a conidia suspension in Roswell Park Memorial Institute medium supplemented with 2% glucose (RPMI 2% G) is injected into the microfluidic channels to introduce the *Aspergillus* conidia to the microwells and given 15 min to allow the conidia to settle within the microtopology (Figure 1A). Subsequently, RPMI 2% G medium adjusted to a specific antifungal agent concentration is slowly introduced into the channels and the fungal response is optically monitored by iPRISM (Fig. 1C). If the conidia germinate and hyphal growth occurs on top of the microwells, the intensity of the reflected light decreases over time (Fig. 1D-ii), while inhibition of growth and cell death result in unchanged intensity values (Fig. 1D-i)

**Figure 1.**
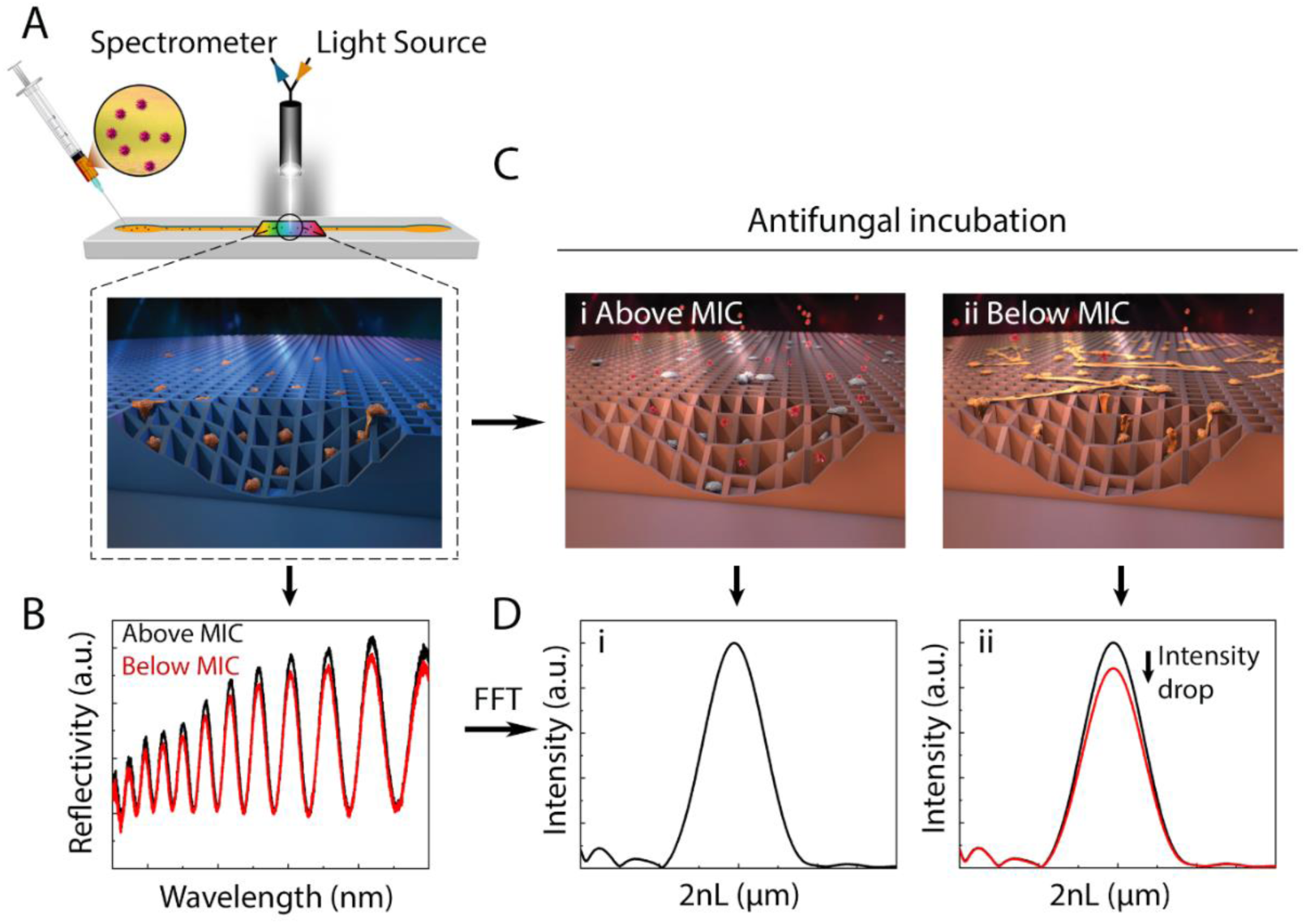
Schematic representation of optical monitoring of *Aspergillus niger* growth and responses to antifungal agents by iPRISM. (A) Si chips of microwell arrays entrap *Aspergillus* conidia from a conidia suspension in RPMI 2% G medium while illuminated by a collimated white light source. (B) The resulting reflectance spectra are recorded and analyzed in real time, allowing for label-free monitoring of fungal growth and responses to antifungal agents. (C) After allowing 15 min for the *Aspergillus* conidia to settle within the Si microwells, RPMI 2% G medium at a designated antifungal agent concentration is introduced, (C-i) resulting in growth inhibition and cell death at concentrations above the MIC or (C-ii) unimpeded growth at subinhibitory antifungal concentrations. (D-i) After applying frequency analysis, growth inhibition corresponds to unchanged intensity values, while (D-ii) fungal growth on top of the microwells results in a reduction of the intensity of the reflected light.

### Optimization of the iPRISM Assay

In order to accelerate AFST of *A. niger*, the initial conidia concentration was optimized (Figure 2). Seeding suspensions at different cell densities were injected into the channels and germination and hyphal growth were verified by optical microscopy after 15 h of incubation (Fig. 2A). Entrapment of conidia inside the well gratings was additionally observed by high resolution scanning electron microscopy (HR-SEM) (Fig. 2B-i). Values of *ΔI (%)* were tracked in real time for conidia concentrations that led to uninhibited germ tube formation (Fig. 2C).

**Figure 2.**
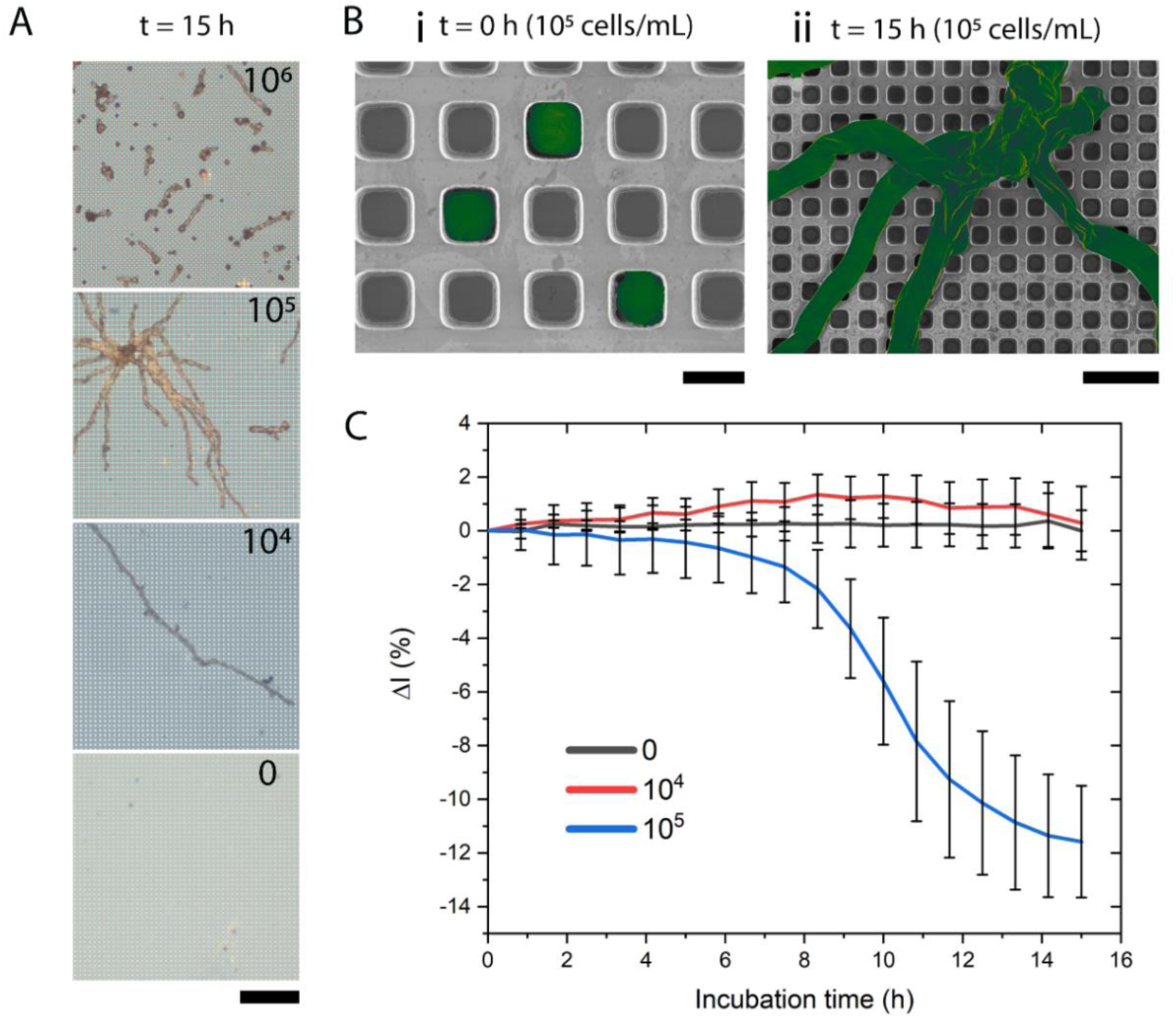
iPRISM for monitoring the growth of *A. niger* at different starting conidia concentrations. (A) Optical microscope images of *A. niger* fixed after 15 h of incubation with conidial seeding suspensions of 0, 10^4^, 10^5^ and 10^6^ conidia mL^−1^. Black scale bar represents 50 µm. (B) HR-SEM images reveal a periodic array of microwells with a diameter of ~3 µm with (B-i) *A. niger* conidia (false-colored green for clarity) entrapped inside the microtopology at t = 0 h (scale bar 3 µm) and (B-ii) *A. niger* spreading over the silicon surface after 15 h of incubation (scale bar 10 µm). (C) *ΔI* (%) over time for conidial seeding suspensions of 0, 10^4^ and 10^5^ conidia mL^−1^ at 25 °C; Average and standard deviation for triplicates (n =3) were calculated every 50 min.

As Fig. 2A demonstrates, the formation of long germ tubes is visible for seeding suspensions with 10^4^ and 10^5^ conidia mL^−1^, while germination and growth are clearly inhibited at an initial conidia concentration of 10^6^ conidia mL^−1^. This inhibition of growth at high conidia concentrations may be due to the fast depletion of nutrients by the high density of cells, or due to self-inhibition.^35^ *A. niger* has been recorded to produce self-inhibitors, such as nonanoic acid^36^, that prevents premature and simultaneous germination of conidia and drastically decrease the germination rate when the cell density is above 10^6^ cells mL^−1^.^37,38^

Corresponding iPRISM curves depict a pronounced decrease in intensity at a seeding suspension of 10^5^ conidia mL^−1^ whereas intensity values for 0 and 10^4^ conidia mL^−1^ are relatively stable. Therefore, we infer that seeding suspensions of 10^5^ conidia mL^−1^ are most suitable to detect and track fungal growth during iPRISM assays for AFST, which corresponds to the suggested seeding concentration by EUCAST protocols for AFST of conidia-forming molds including *Aspergillus* species.^39^ In addition to cell concentration, the temperature of the microfluidic channels during iPRISM assays was optimized as depicted in FigureS1, revealing that changes in intensity occur fastest at 30 °C. Therefore, subsequent experiments were performed at 30 °C and with conidial seeding concentrations of 10^5^ conidia mL^−1^.

### Tracking Aspergillus niger Growth on Well-Type Gratings

To confirm that the decline of intensity corresponds to the growth of *A. niger*, confocal laser scanning microscopy (CLSM) images were acquired after 0, 4, 5, 6 and 15 h of on-chip incubation. *A. niger* conidia and hyphae are stained with calcofluor white (λ_exciation_ = 405 nm) at different time points and the corresponding *ΔI* (%) signal is tracked over time, as demonstrated in Figure 3. During the first 4 h of on-chip incubation, *A. niger* conidia undergo a swelling process as demonstrated by CLSM images (Fig. 3A-ii) and as previously described^40^. However, this swelling process does not result in a corresponding change of the signal intensity (Fig. 3B). Initial germination is apparent after 5 h (Fig. 3A-iii), with further germ tube development and elongation visible after 6 h (Fig. 3A-iv) and 15 h (Fig. 3A-v). The germination of the conidia is accompanied by a slight reduction of the intensity which proceeds and intensifies according to the germ tube elongation; thus, the decline of the signal intensity correlates to the growth of *A. niger*, allowing for the detection of *A. niger* growth using iPRISM.

**Figure 3.**
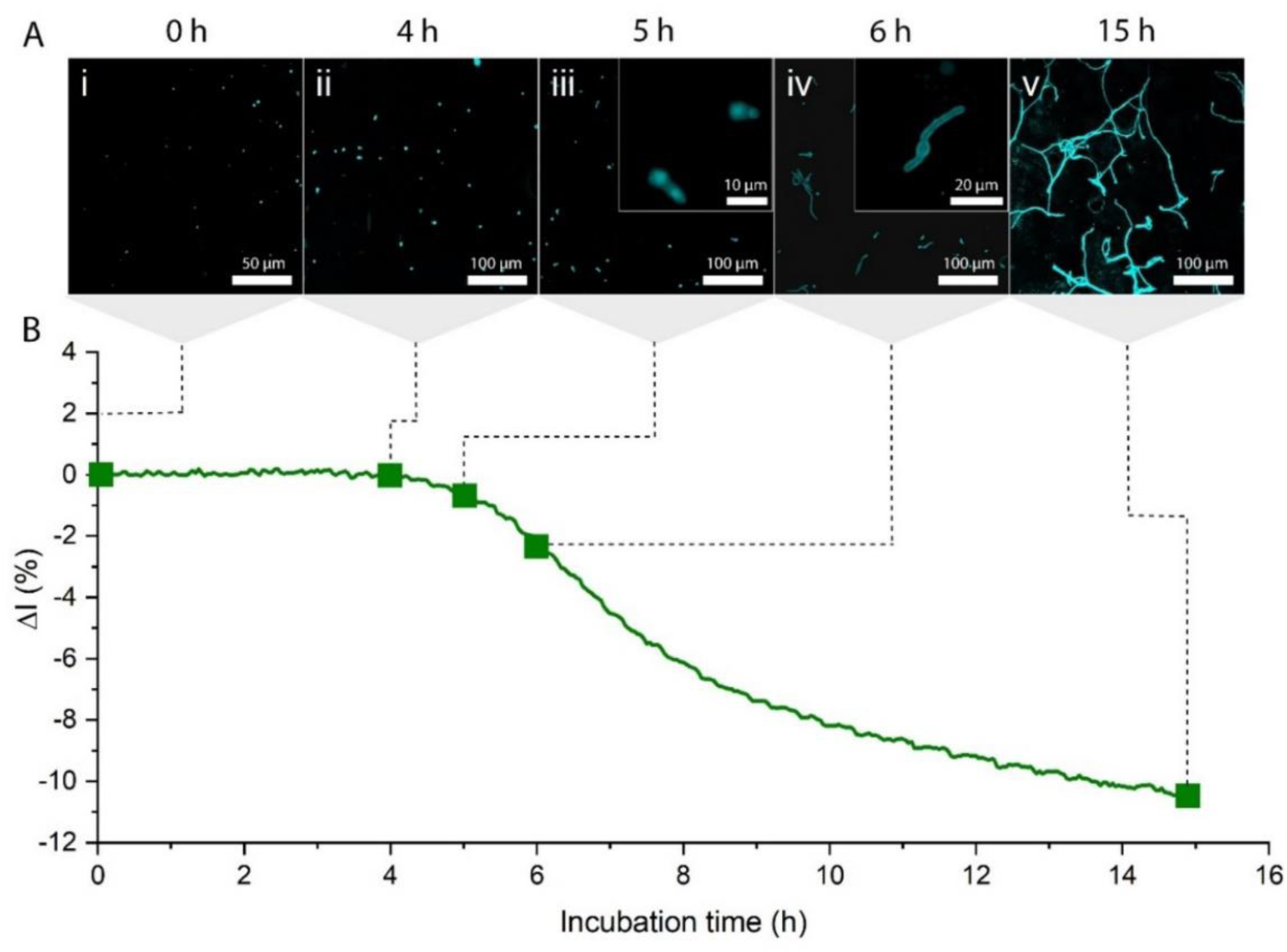
*A. niger* growth on Si microwell gratings at 30 °C and a conidial seeding suspension of 10^5^ conidia mL^−1^. (A) False-colored CLSM images following calcofluor white staining after 0, 4, 5, 6 and 15 h of incubation. (B) Real-time iPRISM curve, where *ΔI* values were recorded over a time period of 15h.

### iPRISM for Antifungal Susceptibility Testing

For the determination of MIC values by iPRISM, *A. niger* conidia were exposed to various concentrations of the clinically relevant antifungal agents voriconazole and amphotericin B (Figure 4). Voriconazole is the recommended antimicrobial agent for treatment of invasive aspergillosis^41^ and inhibits the biosynthesis of ergosterol, which is a crucial component of the fungal cell membrane^42^. Amphotericin B is available for second line treatment and interacts with ergosterol in the cell membrane, resulting in cell death.^43,44^ In the iPRISM assay the MIC is defined as the lowest concentration of an antifungal agent at which no reduction of the intensity value *ΔI (%)* occurs while at subinhibitory concentrations a decline of intensity should be visible to some extent. Moreover, it is expected that the reduction of the signal intensity becomes more prominent with decreasing antifungal concentration.

**Figure 4.**
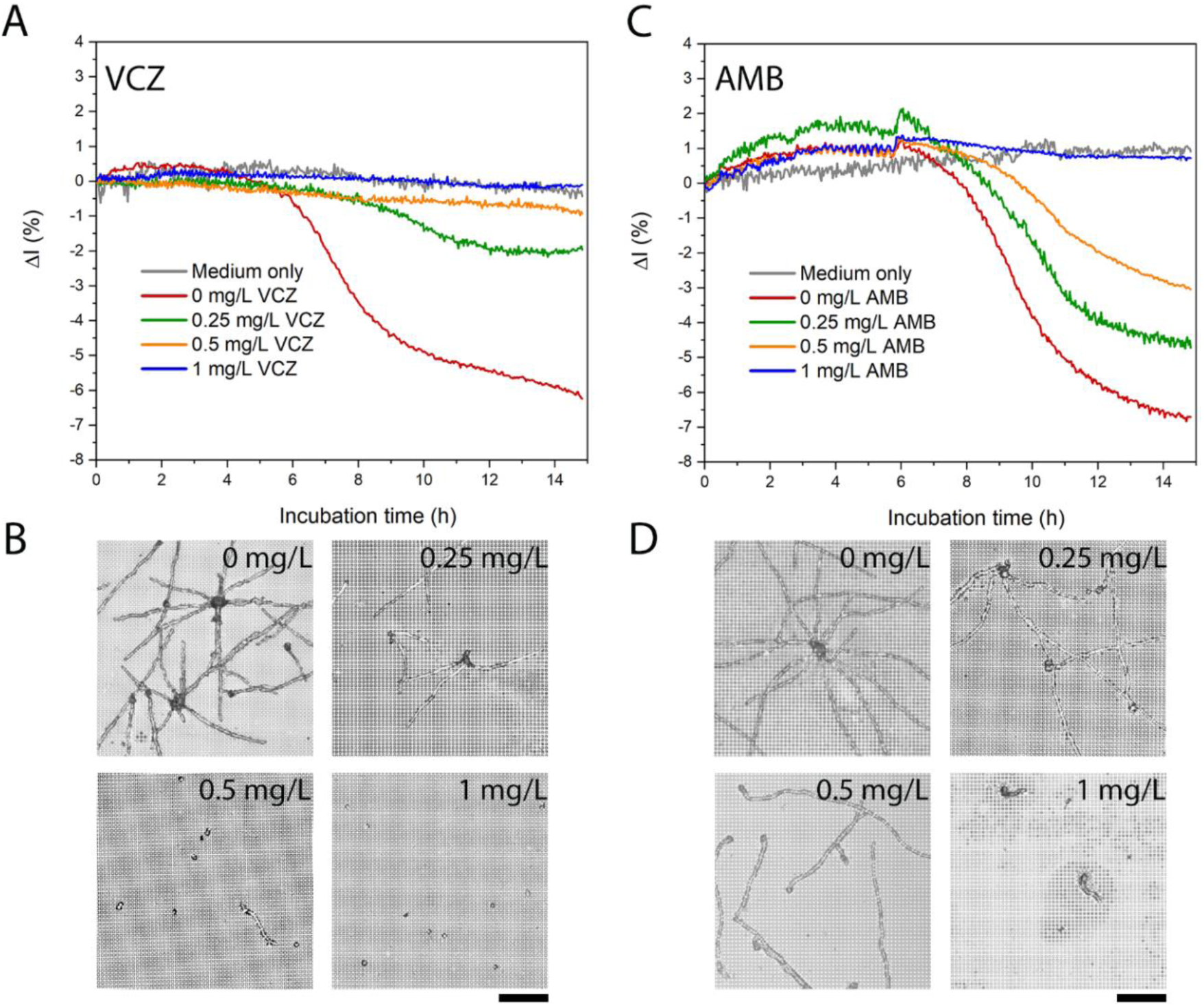
MIC determination of voriconazole (VCZ) and amphotericin B (AMB) against *A. niger* at 30 °C using iPRISM on microwell gratings. (A) *ΔI* (%) over time for various voriconazole concentrations tested against *A. niger*. The MIC was determined to be 0.5 mg L^−1^. (B) Corresponding optical microscope images of chips incubated with various concentrations of voriconazole after 15 h. Scale bar represents 50 µm. (C) *ΔI* (%) plotted versus time for different amphotericin B concentrations revealing a MIC of 1 mg L^−1^ and (D) corresponding optical microscope images. Scale bar represents 50 µm.

Accordingly, with increased voriconazole and amphotericin B concentrations, a decrease in intensity becomes less prominent (Fig. 4A, C) and the growth of *A. niger* is increasingly inhibited (Fig. 4B, D). Following our definition, the MIC for voriconazole was determined to be 0.5 mg L^−1^ and 1 mg L^−1^ for amphotericin B by iPRISM. Microscopy images reveal that at these concentrations the growth of *A. niger* is clearly inhibited (Fig. 4B, D), although minor germination is visible.

Slight variations in the amount of time until a decrease in the intensity is observed and for variations in the outcome of AFST during iPRISM may be due to variation in the inoculum size from assay to assay, particularly as the growth of *A. niger* on the silicon gratings is influenced by cell density. It is known that the inoculum size has an impact on the MIC value.^45,46^ However, comparing the MIC and the highest subinhibitory concentration in triplicate for voriconazole and amphotericin B during iPRISM experiments gives similar results (Figure S2). Moreover, the outcome of AFST by iPRISM was compared to MIC values determined by BMD testing, which is the current gold standard for antimicrobial susceptibility testing (Table 1). The MIC values determined by BMD comply with the MIC distribution for voriconazole and amphotericin B according to the European Society on Antimicrobial Susceptibility Testing.^47^

**Table 1.** MIC determination for voriconazole and amphotericin B on A. niger, evaluated by iPRISM and BMD.

| Antifungal Agent | iPRISM | BMD |
| --- | --- | --- |
| Voriconazole | 0.5 mg/L | 0.5 mg/L |
| Amphotericin B | 1 mg/L | 0.25 mg/L |

The MIC for voriconazole (0.5 mg L^−1^) matches both BMD and iPRISM methods, while the MIC for amphotericin B determined by iPRISM (1 mg L^−1^) is higher compared to BMD (0.25 mg L^−1^). However, it is not surprising that the MIC determined by different methods can vary, as previously the MIC of *E. coli* on silicon gratings was shown to be slightly higher using PRISM than BMD [24]. A main reason for this discrepancy is likely to be due to the different surface structure of our sensor compared to a plain plastic surface and mostly liquid environment of a microwell plate. It is possible that the microstructured well gratings of our silicon sensor also provide a surface that *A. niger* can interact with or perhaps some of the antifungal agent is adsorbed to the silicon surface, increasing the MIC for iPRISM.

## Conclusion

The presented work is a proof-of-concept study, demonstrating the employment of Si diffraction gratings as a label-free assay for real-time antifungal susceptibility testing of filamentous fungi. Differentiation between growth and no growth and determination of MIC values can be achieved within 10 h by iPRISM, thus accelerating the time it takes to generate MIC values compared to classical methods such as BMD or agar-based methods for filamentous fungi. While this platform was demonstrated for *A. niger*, iPRISM can potentially also be used as a tool for monitoring other microorganisms, such as yeast or bacteria due to its simple principle of detection.

## Materials and Methods

### Materials

Glutaraldehyde, Calcofluor white, Roswell Park Memorial Institute medium (RPMI 1640), D-glucose, amphotericin B and voriconazole were supplied by Sigma-Aldrich, Israel. Absolute ethanol, DMSO and all PBS salts were purchased from Merck, Germany. Isopropanol and acetone were supplied by Gadot, Israel. Potato dextrose agar, potato dextrose broth, bacto agar, yeast extract, and casein hydrolysate were purchased from Difco, USA. 3-(*N*-morpholino)propanesulfonic acid (MOPS) was supplied by Chem-Impex International, Inc., USA.

### Preparation of solutions and media

All aqueous solutions and media were prepared in Milli-Q water (18.2 MΩ cm). PBS was composed of 137 mM NaCl, 2.7 mM KCl, 1.8 mM KH_2_PO_4_, and 10 mM Na_2_HPO_4_. PDYC medium was prepared from 24 g L^−1^ potato dextrose broth, 2 g L^−1^ yeast extract and 1.2 g L^−1^ casein hydrolysate. RPMI 1640 2 % G medium was constituted of 10.4 g L^−1^ RPMI 1640, 34.5 g L^−1^ MOPS and 18 g L^−1^ glucose. 1% Potato Dextrose Agar (PDA) contained 10 g L^−1^ PDA and 15 g L^−1^ Bacto Agar, water agar contained 20 g L^−1^ Bacto agar. All media and buffer were autoclaved or sterile filtered prior to use. Filtered stock solutions of amphotericin B and voriconazole were prepared at a stock concentration of 3200 mg L^−1^ in DMSO and diluted in RPMI 1640 2 % G medium prior to iPRISM and BMD testing.

### Aspergillus niger isolate

*Aspergillus niger* was isolated from a contaminated onion into 1% PDA and incubated in darkness for 10 days at 25ºC. Conidia from the mature culture were re-cultured by streaking onto sabouraud dextrose agar plates (SDA) with chloramphenicol (Novamed Ltd., Israel) to eliminate bacterial contamination and incubated as described above. For preparation of mono-conidial cultures, conidia were re-cultured onto water agar and incubated overnight (18h) at 25ºC. Single germinating conidia were removed by micromanipulation under light microscopy and transferred to 1% PDA. All tests in the present study were performed with a single mono-conidial culture isolate designated HCN 18 in order to limit the possibility of genetic variability that may affect tests outcome.

### DNA extraction

*A. niger* isolate HCN 18 was grown overnight at room temperature in liquid PDYC medium in a petri dish without shaking. For DNA extraction two full spatulae of the mycelium were transferred to a clean tube with 100 µL Zirconia/Silica beads (Biospec Products, Inc., USA) and 500 µL DNA extraction buffer (200mM Tris HCL, 0,5 % SDS, 25 mM EDTA, 250 mM NaCl) and mixed for one minute on a vortex mixer (Scientific Industries, Inc., USA). The sample was incubated for 30 min at room temperature and centrifuged for 10 minutes at 4 °C and 12500 rpm (Centrifuge 5417R, Eppendorf AG, Germany). The supernatant was transferred to a clean tube, and an equal volume of isopropanol was added. The sample was incubated for 20 minutes at −20 °C and afterward centrifuged at 13000 rpm for 2 minutes. Subsequently, the supernatant was discarded, and the resulting pellet was washed with 1 mL 70 % ethanol followed by centrifugation at 13000 rpm for 2 minutes. The supernatant was rejected again, and the pellet was left for drying for around 1 h and resuspended in 50 µL of sterile double distilled water. The concentration of the genomic DNA was determined by using a Nanodrop one (Thermo Scientific, USA) resulting in a concentration of 56.4 ng/µL with an A260:A280 ratio of 1.85.

### PCR Amplification of ITS

Species identification of isolate HCN 18 was carried out by sequencing the ITS1 and ITS2 internal transcribed spacer regions by using the primers ITS1F (5’-CTTGGTCATTTAGAGGAAGTAA-3’) and ITS4 (5’-TCCTCCGCTTATTGATATGC-3’).^48^ PCR amplification reaction mixtures contained 1 x PCR buffer, 0.2 mM premixed dNTP’s, 1.25 U of Takara Taq™ Polymerase (all Takara, Japan), 10 µM ITS1F and ITS4 primers and 50 ng genomic DNA in a total volume of 25 µL. For non-template control the PCR reaction mixture contained all the components as mentioned above but no genomic *Aspergillus* DNA. PCR amplification was carried out in a biometra TGradient Thermocycler (Analytik Jena, Germany) using an initial denaturation step at 94 °C for 2 min, followed by 29 cycles at 94 °C for 30 s, 55 °C for 30 s and 72 °C for 30 s, with a final extension for 2 min at 72 °C. Successful amplification was confirmed after standard electrophoresis in a 1% agarose gel by visualizing the PCR product under UV light by using the SimplySafe™ dye (EURx, Poland). The PCR fragment was extracted from the gel by using a DNA gel extraction kit (AccuPrep Gel Purification Kit, Bioneer, South Korea) and sent to Hylabs (Israel) for sequencing by using an ABI 3730xl DNA Analyzer (Thermo Scientific, USA). The sequencing result (Table S1) was investigated in BLASTn (https://blast.ncbi.nlm.nih.gov/Blast.cgi?PAGE_TYPE=BlastSearch) with default search parameters and identified the isolated fungus as *Aspergillus niger* (NCBI Accession number: MK182796.1, https://www.ncbi.nlm.nih.gov/nuccore/MK182796)

### Preparation of Fungal Cultures

Fungal cultures were refreshed every two weeks onto 1% PDA and maintained at 4 °C until used. Prior to iPRISM experiments the cultures were transferred onto SDA plates containing chloramphenicol, to eliminate potential bacterial contamination. Because the *Aspergillus* showed the fastest growth at 30 °C, the fungal cultures were subsequently incubated at 30 °C and after 2 – 5 days sufficient sporulation for iPRISM and BMD was reached. The conidia were gently removed from the agar plate with a sterile cotton swab and resuspended in sterile double distilled water. The suspension was centrifuged for 20 min at 5000 rpm (Sigma 2-16P, Sigma GmbH, Germany) and the pellet was resuspended in sterile filtered RPMI 2% G medium. Conidial density was quantified using the aid of a hemocytometer (Neubauer improved cell counting chamber), and the respective conidial densities for iPRISM and BMD were obtained by dilutions in RPMI 2% G medium.

### Fabrication and preparation of photonic silicon chips

Si microwell gratings with a diameter of ~ 3 µm and a pore length of ~ 4 µm were fabricated by standard lithography and reactive ion etching techniques at the Micro- and Nano-Fabrication and Printing Unit (MNFPU, Technion – Israel Institute of Technology). The resulting wafers were coated with photoresist to protect the microstructure while dicing the wafer into 5 x 5 mm chips using an automated dicing saw (DAD3350, Disco, Japan). The silicon chips were washed with acetone to remove the photoresist and oxidized for 1 h at 800 °C in a furnace (Lindberg/Blue M 1200 °C Split-Hinge, Thermo Scientific, USA).

### iPRISM Assay

A custom-made, aluminum flow chamber with seven injection and outlet channels was used to fix and separate the photonic silicon chips during the iPRISM assays. Each injection channel was connected by tubing to a syringe injection port and allowed for the injection of the conidia suspensions. The chamber was controlled by a motorized linear stage (Thorlabs, Inc, USA). The photonic chips were placed in a small square cavity in each channel and were separated from each other and fixed on the surface of the chamber by a rubber gasket. The system was further sealed before the experiments by an acrylic piece and by tightening the lid of the aluminum housing on the acrylic spacer and the rubber gasket. Before each experiment, the system was sterilized by washing with 70% ethanol and sterile water followed by the injection of 500 µl RPMI 2% G medium to allow devices, temperature, and medium to equilibrate. 500 µL of the conidia suspension were slowly injected while the reflectance signal was recorded continuously during the experiment.

### Data acquisition and analysis

A bifurcated fiber optic (Ocean Optics, USA) equipped with a collimating lens was positioned normal to the photonic silicon chips, illuminating them via a white light source. The reflected light was recorded by a USB4000 CCD spectrometer (Ocean Optics, USA). The position of the aluminum chamber was controlled by a motorized linear stage (Thorlabs) and LabView (National Instruments, USA). Frequency analysis was perfomed on acquired spectra in a range between 450 to 900 nm. The resulting peak after fast Fourier transform (FFT) was identified by determining the maximum peak position where the height of the detected peak corresponds directly to the intensity of the reflected light. The intensity values were plotted versus time and the moment when the conidia suspension was injected into the system was defined as time 0 during growth monitoring experiments. For antifungal susceptibility testing the moment of injection of the antifungal agent was instead defined as time 0. The percent changes of the intensity (ΔI) was calculated as follows:

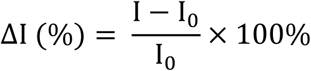

### Characterization of Silicon Chips

The chips were examined immediately after experiments by using an optical light microscope (Axio Scope A1, Carl Zeiss, Germany) to identify fungal growth and assure no bacterial contamination occurred.

High-resolution scanning electron microscopy (HR-SEM) was performed using a Zeiss Ultra Plus high-resolution scanning electron microscope fitted with a Schottky field-emission gun (Carl Zeiss, Germany). The silicon chips were prepared for microscopy analysis by fixation of the chips with 2.5% glutaraldehyde in PBS, followed by a washing step in water and dehydration through a dilution series in ethanol with increasing concentration from 10% to absolute ethanol. Subsequently, the photonic chips were dried gently under a stream of nitrogen.

Confocal laser scanning microscopy (CLSM) was performed on samples stained by Calcofluor white and a drop of 10% potassium hydroxide for better visualization, by using an LSM 700 laser scanning microscope (Carl Zeiss, Germany). The fluorescent dye was excited at (λ_exciation_ = 405 nm) and the images were rendered by Zen software (Carl Zeiss, Germany).

### Broth Microdilution

Broth microdilution was performed according to the EUCAST protocol for antifungal susceptibility testing against conidia forming molds.^47^ Two-fold serial dilutions of the antifungal agents were made in RPMI 2% G medium. The effect on the fungal growth in a conidia suspension with 10^5^ conidia mL^−1^ was observed after 48 h visually and by OD_600_ measurements (n = 5, Varioskan Flash, Thermo Scientific, USA). One adjustment was made compared to the EUCAST protocol: The samples were incubated for two days at 30 °C instead of 35 °C to 37 °C because the iPRISM assays for antifungal susceptibility testing were also performed at 30 °C.

## Associated Content

### Supporting Information

The following files are available free of charge and included in the Supporting Information: Temperature influence experiments, replicate data experiments, and genomic sequencing results.

## Author Information

### Notes

The authors declare no competing financial interests.

## Acknowledgements

We acknowledge the financial aid from the PROMOS scholarship and the Leibniz Universitätsgesellschaft for assisting C.H. in his graduate studies and Joan and Irwin Jacobs for assisting H.L in her doctoral fellowship. The authors also thank the Institute Merieux, Israeli Ministry of Science, and the Nesta Discovery Award for financial assistance. Furthermore, we thank Dima Peselev and Orna Ternyak (MNFPU, Technion – Israel Institute of Technology) for the microfabrication of the photonic silicon chips and Andy Henik for his artwork of *Aspergillus niger* interacting with the silicon well gratings, as well as Dima Abelski for his artwork depicting the microfluidic channels.

