## Supporting Information for "Antifungal susceptibility testing of *Aspergillus niger* on silicon microwells by intensity-based reflectometric interference spectroscopy"

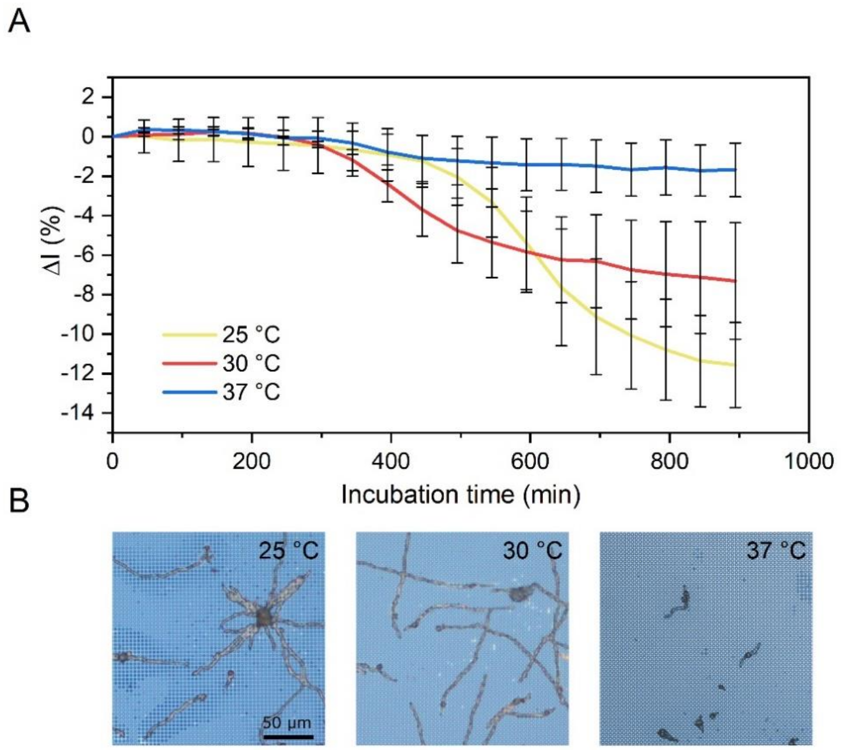


**Figure S1.** Influence of the temperature on *A. niger* growth and detection via iPRISM. (A) $\Delta I$ (%) over time for different temperatures (25, 30, 37 °C) with a seeding suspension of $10^5$ conidia mL$^{-1}$; Average and standard deviation for triplicates (n =3) were calculated every 50 min. (B) Corresponding optical microscope images after 900 min; Scale bar represents 50 µm.

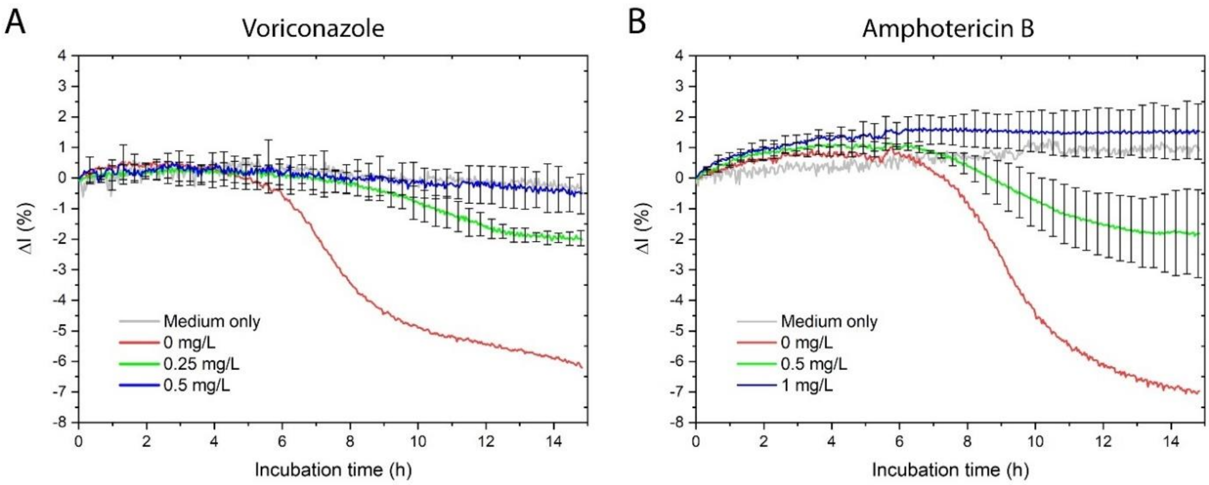


**Figure S2.** Comparison of the MIC and the highest subinhibitory concentration of voriconazole and amphotericin B against *Aspergillus niger* in triplicate (n =3). (A) Comparison of the MIC (0.5 mg $L^{-1}$) and highest subinhibitory concentration (0.25 mg $L^{-1}$) of voriconazole against *A. niger* according to Figure 4. (B) Comparison of MIC (1 mg $L^{-1}$) and highest subinhibitory concentration (0.5 mg $L^{-1}$) of amphotericin B against *A. niger* according to Figure 5.

**Table S1**. Sequencing result for the forward primer ITSF1

| **DNA sequence** |
| --- |
| 5' -<br>TTCCGTAGGTGAACCTGCGGNGGATCATTACCGAGTGCGGGTCCTTTGGGCCCAACCTCCCATCCGTG<br><br>TCTATTGTACCCTGTTGCTTCGGCGGGCCCGCCGCTTGTCGGCCGCCGGGGGGGCGCCTCTGCCCCCCGGGCCCGTGCCC<br><br>GCCGGAGACCCCAACACGAACACTGTCTGAAAGCGTGCAGTCTGAGTTGATTGAATGCAATCAGTTAAAACTTTCAACAA<br><br>TGGATCTCTTGGTTCCGGCATCGATGAAGAACGCAGCGAAATGCGATAACTAATGTGAATTGCAGAATTCAGTGAATCAT<br><br>CGAGTCTTTGAACGCACATTGCGCCCCCTGGTATTCCGGGGGGCATGCCTGTCCGAGCGTCATTGCTGCCCTCAAGCCCG<br><br>GCTTGTGTGTTGGGTCGCCGTCCCCCTCTCCGGGGGGACGGGCCCGAAAGGCAGCGGCGGCACCGCGTCCGATCCTCGAG<br><br>CGTATGGGGCTTTGTCACATGCTCTGTAGGATTGGCCGGCGCCTGCCGACGTTTTCCAACCATTCTTTCCAGGTTGACCT |

CGGATCAGGTAGGGATACCCGCTGAACTTAAGCATATCAATAAGCGGGAGGA
A - 3'
